# Rhizonet: Image Segmentation for Plant Root in Hydroponic Ecosystem

**DOI:** 10.1101/2023.11.20.565580

**Authors:** Daniela Ushizima, Zineb Sordo, Peter Andeer, James Sethian, Trent Northen

## Abstract

Digital cameras have the ability to capture daily images of plant roots, allowing for the estimation of root biomass. However, the complexities of root structures and noisy image backgrounds pose challenges for advanced phenotyping. Manual segmentation methods are laborious and prone to errors, which hinders experiments involving several plants. This paper introduces Rhizonet, a supervised deep learning approach for semantic segmentation of plant root images. Rhizonet harnesses a Residual U-Net backbone to enhance prediction accuracy, incorporating a convex hull operation to precisely outline the largest connected component. The primary objective is to accurately segment the biomass of the roots and analyze their growth over time. The input data comprises color images of various plant samples within a hydroponic environment known as EcoFAB, subject to specific nutrition treatments. Validation tests demonstrate the robust generalization of the model across experiments. This research pioneers advances in root segmentation and phenotype analysis by standardizing processes and facilitating the analysis of thousands of images while reducing subjectivity. The proposed root segmentation algorithms contribute significantly to the precise assessment of the dynamics of root growth under diverse plant conditions.

## 1 Introduction

Biofuels represent renewable energy sources extracted from organic materials, such as plants or plant remnants, and can serve as an eco-friendly substitute for conventional fossil fuels^1^,^2^. Specifically, microbiomes play a pivotal role in nutrient cycling and the promotion of plant growth^3^, both of which are vital to the efficiency of bio-energy crop cultivation. Gaining insight into the plant roots can support research on the optimization of nutrient accessibility, improved nutrient absorption, and enhanced plant growth and biomass yield. However, examining root morphology in soil presents challenges due to its opacity, while investigating excavated root systems hinders the ability to collect root information over time. Moreover, achieving reproducibility across different laboratories adds another layer of complexity to these challenges.

To monitor plants growing under different concentrations of nutrients and environmental conditions, a modular growth system called EcoFAB (Ecosystem Fabrication)^4^ was created. Depicted in Fig. 1, the EcoFAB enables hydroponic, controlled, and reproducible model ecosystems in which microorganisms and host responses can be monitored as a reaction to changing variables^5^. One of the pressing needs is to detect plant roots and quantify their biomass using automated image segmentation^6–8^.

**Figure 1.**
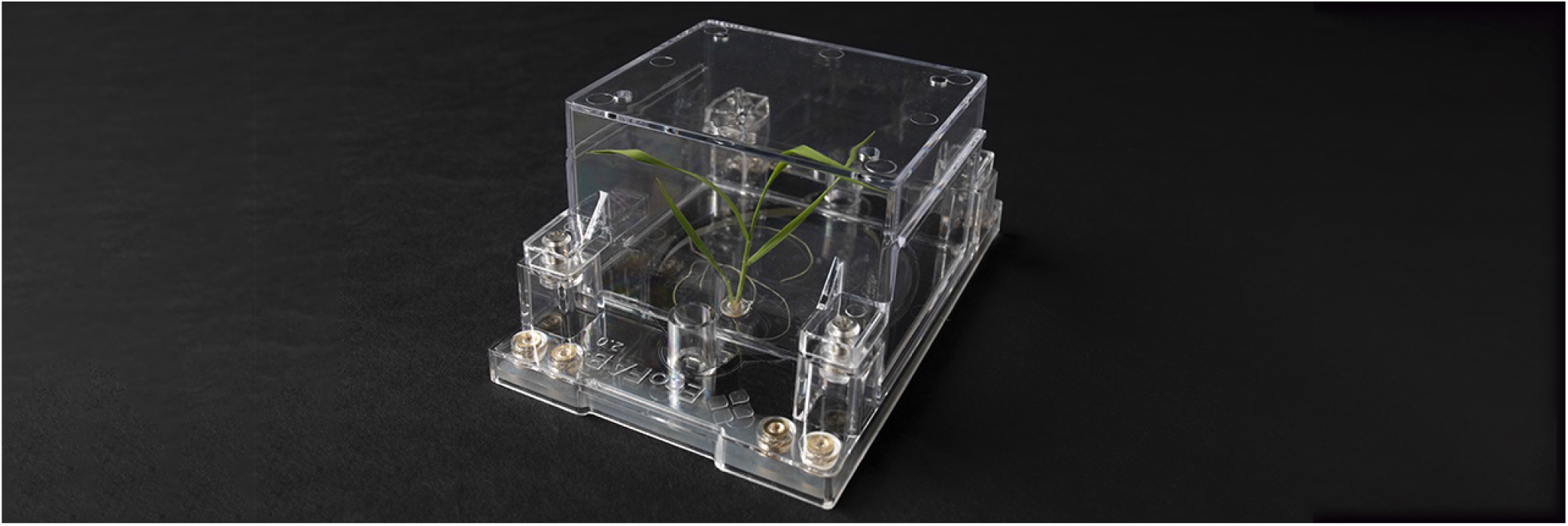
EcoFAB: creating controlled hydroponic model ecosystems for monitoring microbial and host responses to variable environments. Image courtesy: Thor Swift.

Semantic segmentation techniques, particularly those built on deep learning encoder-decoder architectures, have exhibited remarkable results in a variety of scientific fields^9,10^. Recently, these algorithms have been successfully applied to detect abnormal protein clusters present in human brain data from experiments in Alzheimer’s research^11^. Building on these advancements, an enhanced semantic segmentation approach has been employed to detect and evaluate lithium agglomeration in batteries during charge and discharge cycles^12^. Given these diverse applications, could we potentially generalize the algorithm to segment plant roots?

In the field of neuroscience, semantic segmentation methodologies such as IHCNet^11^ applied to immunohistopathology samples have played a pivotal role in the detection and validation of new radioactive tags for Alzheimer’s disease. Similarly, in materials science and battery research, the batteryNet^12^ framework, which uses microCT and 3D + t data, has proven invaluable for identifying lithium agglomeration and evaluating electrolyte quality during battery operation. These accomplishments have deepened our understanding of branching patterns in various contexts, including neurons and battery filaments. This paper introduces a novel approach aimed at adapting and improving our previous efforts and showing the portability of semantic segmentation algorithms across domains, specifically for plant root analysis in EcoFABs.

This paper presents key contributions. In Section 2, we review existing research on segmentation and the use of neural networks for plant images. Section 3 unveils our high-resolution EcoFABs root dataset. We then introduce a novel method to assess root biomass using a convolutional neural network allied to convex hull operators. Our findings are summarized in Section 4- 5 with insights into future automated root segmentation. The workflow is detailed in Fig. 2, showcasing our software’s modularity and segmentation steps.

**Figure 2.**
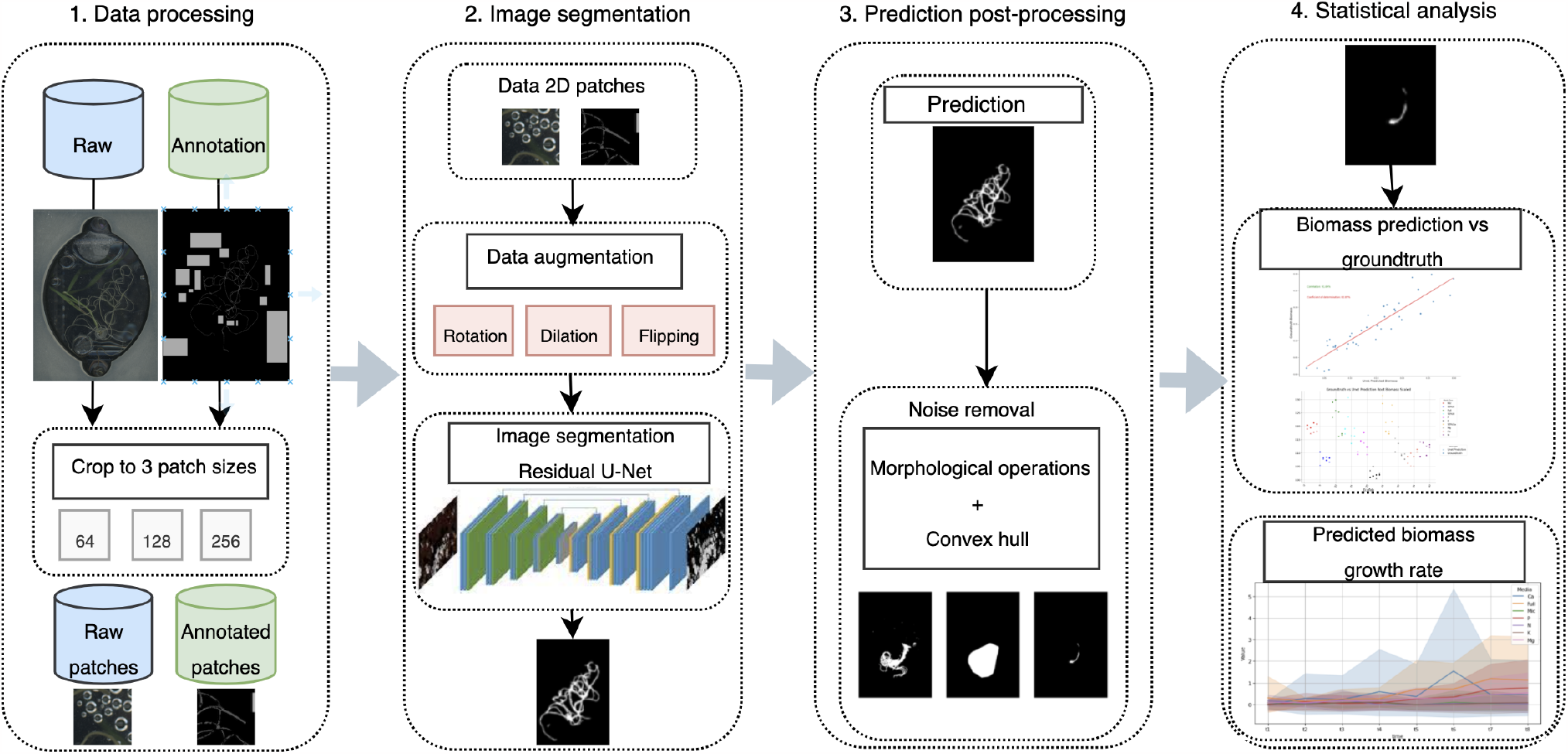
Proposed pipeline for root segmentation and root biomass estimation

### 2 Related work

Root systems play a central role in plant development and nutrient acquisition. To observe root growth and interactions with soil, researchers have often relied on specialized instruments known as minirhizotrons^13^. With early versions dating the 1970s^14^, current minirhizotrons consist of transparent tubes or chambers that are vertically inserted into the soil, enabling the nondestructive monitoring and capture of images of plant roots as they evolve in their natural environment. While minirhizotrons provide valuable insights, they possess limitations such as restricted depth observation and interference with nearby roots. In addition, they require complex data analysis and are best suited for transparent soils. To address these limitations and explore alternative methods of real-time root observation, our study turns to EcoFAB systems, offering a promising alternative for plant surveillance under various nutrient conditions.

In terms of root analysis, threshold-based methods have shown effectiveness in numerous applications^15–17^, particularly when there exists a substantial contrast between the foreground and background, often achieved by sample manipulation, for example, staining techniques, or controlled intensity adjustments. However, grayscale thresholding proved to be inadequate for root segmentation under the unique conditions posed by EcoFABs, where artifacts such as condensation and the presence of delicate, shadow-casting leaves can obscure root visibility. As a result, researchers have turned to U-Net-based approaches^10,11,18^, which have shown remarkable performance in various plant classification tasks^19–21^. Nevertheless, U-Nets may encounter limitations when dealing with intricate, fine structures, such as those encountered in high-resolution EcoFAB imagery. Given the labyrinthine and dynamic nature of root systems within EcoFABs, a more robust and customized solution becomes imperative.

To address these challenges, we introduce RhizoNet, an algorithm that leverages a Residual U-Net architecture combined with convex-hull to deliver root segmentation specifically adapted for EcoFAB conditions. Our approach bridges the gap between traditional root analysis methods and the unique demands of high-resolution images from controlled studies of nutrient deprivation within EcoFABs, enabling more precise and informative insights into root growth and plant responses.

## 3 Materials and Methods

### 3.1 Grass as a research model

*Brachypodium distachyon* is a small grass species that serves as a cost-effective research model to study plants^22,23^. With a small genome size, a short life cycle, and genetic similarity to important cereal crops such as wheat, barley, and oats, it enables the investigation of various aspects of plant biology. Although it is not a major biofuel crop, insights from *B. distachyon* research can inform the development of more efficient biofuel crops by identifying genes and traits related to biomass yield, nutrient utilization, and stress tolerance.

Figure 2 illustrates the computational pipeline for processing roots of *B. distachyon* plants that are subjected to various conditions of deprivation of nutrients for approximately 3 weeks. Each plant grows in its respective EcoFAB chamber, which is monitored for days, with images acquired using a high-resolution flatbed scanner (Fig. 3), providing information on the evolution of the plant over time within an EcoFAB.

**Figure 3.**
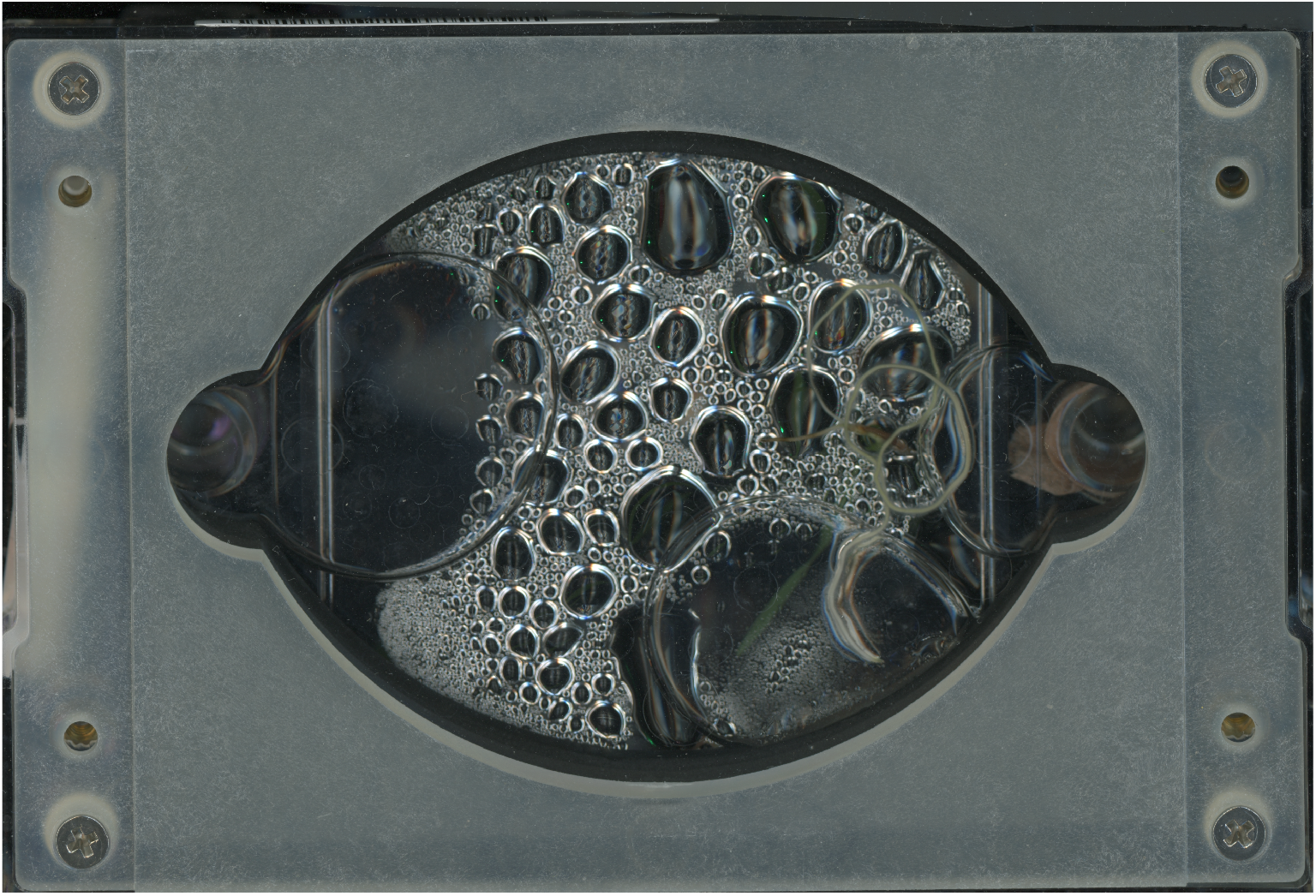
Non-processed EcoFAB image after two weeks: acquired on September 1st 2022, with visible condensation.

EcoFABs represent artificial microbial ecosystems, offering standardized and reproducible model systems for the investigation of microbiomes^24^. EcoFABs comprise several essential components:

- Field Site: A reference ecosystem, providing soil, microbes, and a place for long-term validation of findings.
- Model Plant: Well-characterized plants, such as *Brachypodium distachyon*, are genetically transformable and can complete their life cycle within EcoFab.
- Model Bacteria: Bacteria sources from the field site, both isolated and intact communities.
- Model Soil: Ceramic substrates or synthetic soils provide a stable, reproducible, and sterilizable growth medium.
- Growth Chambers: Controlled environments with flexible designs, potentially using 3D printing technology, to facilitate plant growth, microbial interactions, and observations.

To capture images of plant roots, the lower section of each EcoFAB undergoes a scanning process, as depicted in Fig. 3 and detailed in the next section.

### 3.2 Data acquisition and inherent artifacts

For image acquisition, we used a high-end scanning device, the EPSON Perfection V850 Pro. This professional-grade scanner is designed for advanced scanning applications, offering exceptional precision and quality when scanning paper sheets and various other media types. The key features of the EPSON Perfection V850 Pro include a versatile scanning solution with its dual high-resolution lens system, capable of scanning photos, documents, and 35mm film and slides. It boasts an impressive optical resolution of up to 6400 dpi, enabling detailed and sharp scans at fast scanning speeds. Advanced scanning technologies, including the Dual Lens System, Micro Step Drive, and Digital ICE™for dust and scratch removal, ensure high-quality and accurate scans. Furthermore, its wide compatibility with both Windows and Mac operating systems makes it accessible to a diverse range of users.

Despite our ability to achieve high-quality scans, the images of plant roots generated during the EcoFab-based plant growth process present intricate challenges for root detection. This process introduces various issues, including condensation, bubbles, and shadows. Condensation in plant imaging arises from the formation of water droplets on surfaces of imaging equipment due to temperature disparities between the equipment and its environment. This frequently occurs in environments with temperature variations, such as growth chambers, where warm, humid air contacts cooler surfaces. The condensation issue becomes particularly problematic during extended imaging sessions, notably in the later frames of the time series (refer to Fig.3). Similarly, but on a larger scale, bubbles can form due to plant roots, primarily through oxygen release, which benefits the microorganisms in the root zone but can obstruct plant root visibility in the proposed imaging system.

These physical phenomena pose difficulties for segmentation algorithms, which may misinterpret subtle droplet edges as roots. Furthermore, leaf shadows also introduce background noise, complicating the distinction between thin leaves and the target for both algorithms and human observers due to their pixel-wise resemblance to roots.

To mitigate these artifacts, two experiments were carried out, namely `Exp 1` and `Exp 2`, using a *B. distachyon* seedling per EcoFAB and dozens of EcoFABs per experiment, as described in Table 1.

**Table 1.** Distribution of plant root scans in EcoFABs acquired during different experiments: Exp = experiment, Nb = number.

| Exp | Nb of EcoFABs | Nb of scans | Nb of training scans | Nb of val/test scans | Nb of unseen scans |
| --- | --- | --- | --- | --- | --- |
| 1 | 40 | 337 | N/A | N/A | 337 |
| 2 | 42 | 306 | 61 | 10 | 245 |

### 3.3 Dataset composition and experimental conditions

The dataset consists of images acquired from two separate experiments conducted in various environmental settings. Each experiment featured distinct EcoFab plants exposed to specific nutritional conditions, including variations in nitrogen, calcium, magnesium, potassium, and micronutrient concentrations (refer to Fig. 4). Eight images were captured for each EcoFab plant, covering eight different dates with intervals ranging from 3 to 7 days.

**Figure 4.**
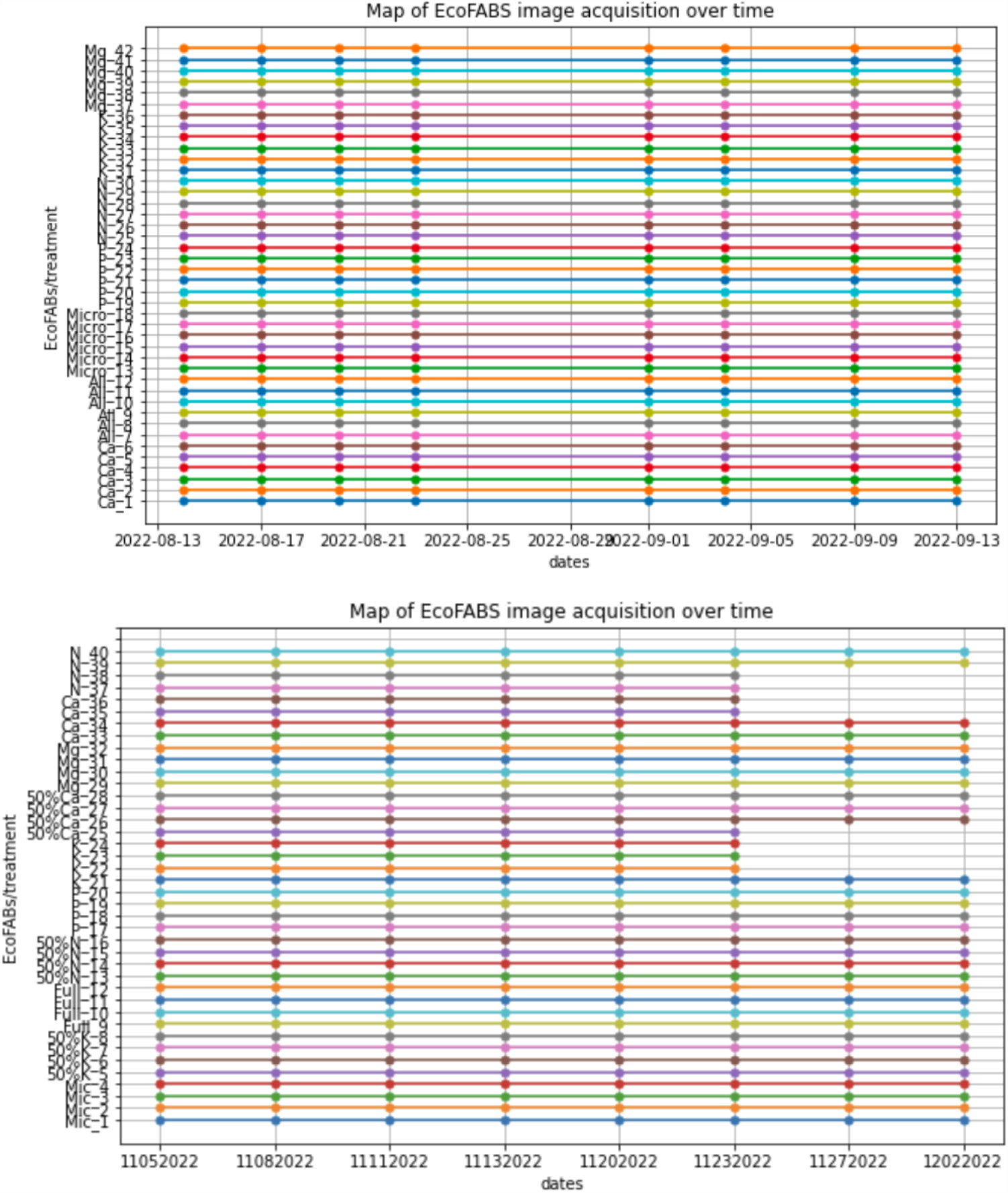
Timeseries of images in Experiment 1 and 2 (bottom), representing each EcoFab scan as a circle.

In Experiment 2, conducted in an environment with a lower thermostat setting, two notable aspects emerged. Firstly, this setup provided an advantage by enabling the acquisition of superior images of the root system at reduced temperatures, minimizing issues related to condensation and bubble formation. However, a significant disadvantage emerged as plants subjected to potassium and calcium deficiencies ceased growth and eventually perished after a few days. This particular challenge was unique to Experiment 2 and was attributed to the colder conditions compared to Experiment 1, where all plants exhibited continuous growth without evident necrosis until the final observation date.

### 3.4 Image annotation

For data preparation, we initiated the process by identifying and labeling three distinct classes: background (class 0), noise (comprising droplets, bubbles, or leaves, labeled class 1) and root (class 2). As highlighted in Table 1, only a subset of the second dataset underwent annotation, therefore only this subset was subsequently used for training, validation and testing of the segmentation algorithms. The unlabeled data served the purpose of making additional predictions using the weights of the trained model, allowing us to evaluate its performance on previously unseen data.

The groundtruth biomass measurements were conducted using a scale (balance) to measure root weights for both experiments. However, these measurements were performed exclusively on the last acquisition date for each EcoFab. Consequently, this partial groundtruth corresponds to the biomass recorded on the final date for each of the EcoFabs measured.

In contrast, the biomass of the manually annotated images was calculated by quantifying the number of pixels corresponding to the root. Subsequently, this value was normalized, as was the groundtruth biomass, to facilitate comparative and analytical evaluations. It is worth acknowledging that some degree of uncertainty may be present in individual root annotations, which should be considered in further interpretations.

Our analysis commenced with an evaluation of the biomass of the manual annotations concerning the ground-truth biomass (balance) of the final acquisition date within the second dataset. We performed a regression analysis, initially fitting a model between the labeled images and the groundtruth. This approach provided correlation values, including specific correlations to different nutritional deficiencies (Table 2), further discussed in the Results section (refer to Fig. 10).

**Table 2.** Correlation values for each nutrition condition observed in Exp 02.

|  | Nutrition Deficiency | Pearson correlation |
| --- | --- | --- |
| 1 | 50%Ca | 93.6% |
| 2 | 50%K | 95.1% |
| 3 | 50%N | 89.6% |
| 4 | Full | 91.8% |
| 5 | Mg | 94.8% |
| 6 | Mic | 14.3% |
| 7 | N | 89.6% |
| 8 | P | 70% |
| 9 | Ca | 71.2% |

## 3.5 Data analysis

The images annotated by the human experts compose the training dataset, consisting of 61 original scans of size (3000, 2039) extracted from the sixth and eighth acquisition dates of Exp 2, and 61 grayscale annotated target images (Fig. 5). Given that the raw data contain noise and include irrelevant parts due to condensation, bubbles, and shadows, we proceed to develop an algorithm that includes routines to ignore those regions.

**Figure 5.**
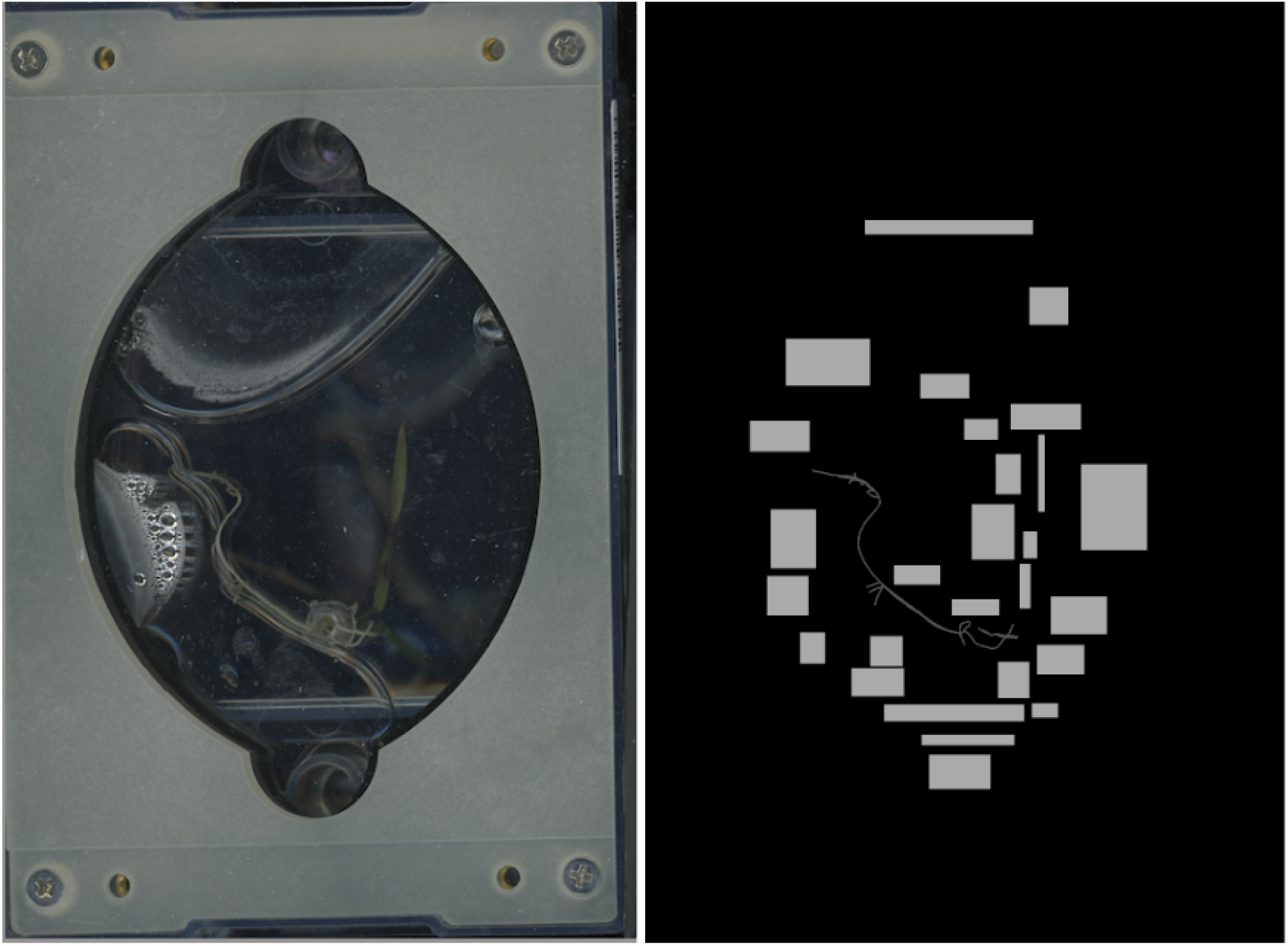
Full size sample: *left*: raw input image with dimensions (3000, 2039) pixels and *right*: corresponding labeled image as annotated by a human expert.

Therefore, to enable the algorithm to focus on the lower-level features of the root and better understand its structure, we created three different types of patches of sizes (64,64), (128,128) and (256,256). We considered 2D models capable of handling RGB images and subdivided each training hand-labeled slice and raw image into 128 × 128 patches (refer to Fig. 6), 64 × 64 patches and 256 *×* 256 patches, which were then separated into 80% training, 10% validation and 10% testing datasets. This processing was especially helpful given that each image was highly imbalanced pixel-wise since the very thin roots occupied a very small part of the (3000, 2039) raw images.

**Figure 6.**
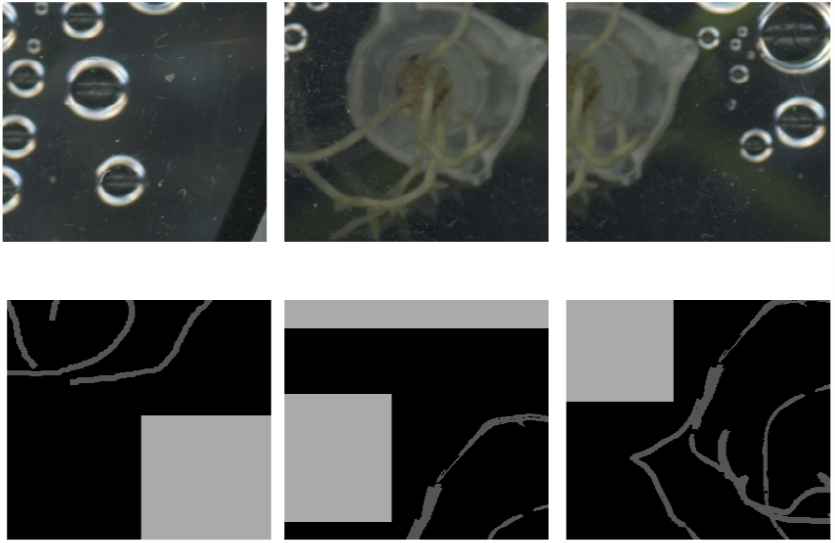
Training patch sample: *top*: random raw input patches of size (128,128) and *bottom*: random labeled patches of size (128,128)

In addition to creating a training dataset of patches, we also executed data augmentation schemes in training all models, which consist of random rotations in all directions, random flips (vertically and horizontally), random cropping (2%), random shifts, random zoom (range in [0.8, 1]), and a small range of random brightness and contrast variation (*±* 5%). These geometric transformations are applied at run-time before the training of the RhizoNet begins. This data augmentation aims at diversifying the training dataset and reducing overfitting in the model.

### 3.6 Computational models

In developing our software tool, we employ the scientific Python software stack, encompassing key libraries like scikit-learn for machine learning and matplotlib and seaborn for visualization. We also integrate *MONAI* (Medical Open Network for AI) — an open-source, *PyTorch*-based framework tailored for healthcare imaging. *MONAI* offers specialized functionalities to develop deep learning techniques for biological imaging tasks^25,26^ and optimizes the code for rapid processing. Crucially, it facilitates GPU acceleration in our algorithm, significantly boosting the speed of tasks like deep learning-based image segmentation. This enhancement allows our software to swiftly handle large image data sets, aligning with the rigorous requirements of plant scientific research.

The creation of the RhizoNet model benefits from high performance resources at NERSC, by using a node of the system called Perlmutter. It enables accelerated training using the NVIDIA Tesla V100 GPU for each experiment, the model size was roughly 26 MB for 6.5 million parameters. The following Table 3 summarizes the number of patch samples for each training and the associated duration of training.

**Table 3.** Patch sizes, number of patches used during training and training time for creation of RhizoNet models - sample size before augmentation.

| Patch Size | Sample size | Training time |
| --- | --- | --- |
| (64,64) | 38176 | 30 min |
| (128,128) | 10943 | 45 min |
| (256,256) | 3209 | 1 hour |

### 3.7 Deep Learning: Residual U-Net

Figure 7 shows a convolutional neural network known as U-Net, an architecture that was first introduced in 2015 by Ronneberger et al. for the semantic segmentation of biomedical images. The original work^9^ proposes an architecture that consists of a contracting path to capture context and a symmetric expanding path that enables precise localization. In our work, the model was adapted to take images of size 128 *×* 128 as input. For the encoder, we started with 32-channel 3 *×* 3 kernels but each standard block was replaced by a 2-convolution residual unit, which is detailed in the next section.

**Figure 7.**
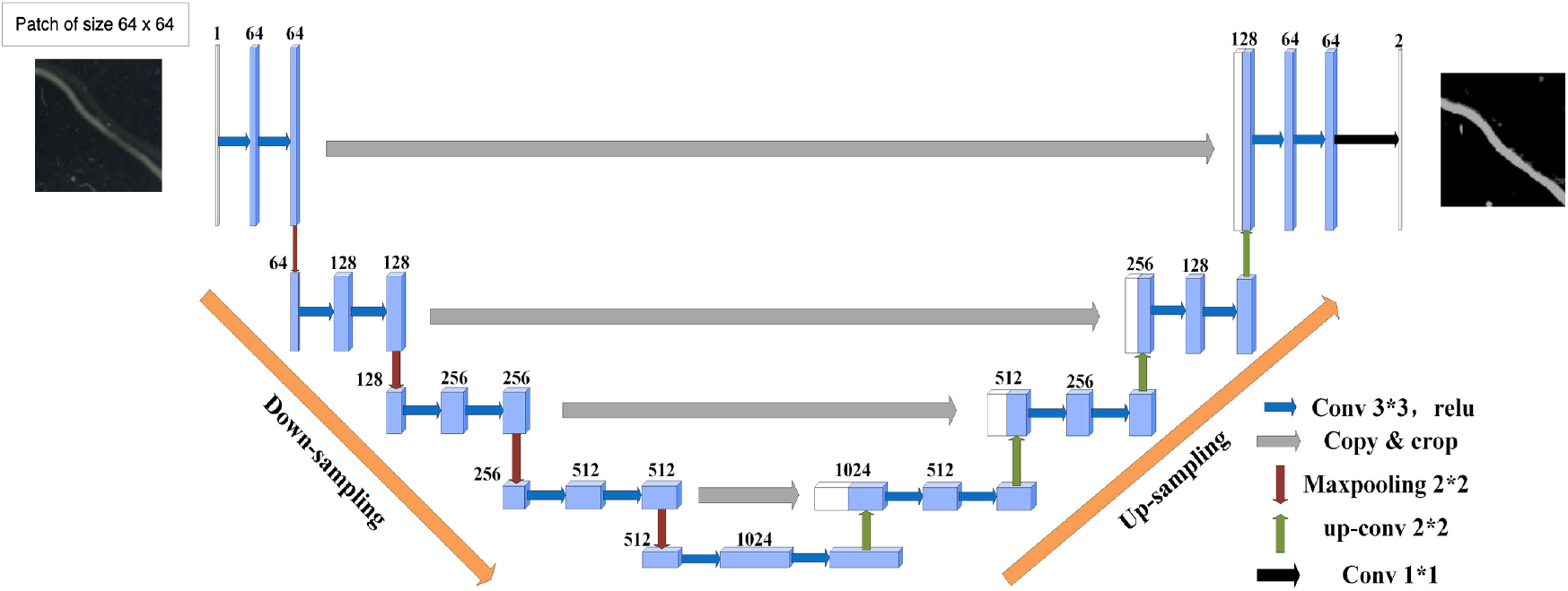
U-Net architecture for (64,64) input images with 5 layers in both Encoder: 5 Convolutional Layers (3×3, ReLU Activation), Max-Pooling (2×2) and Decoder: 5 Convolutional Layers (3×3, ReLU Activation), Max-Pooling (2×2).

An enhanced version of U-Net is the Deep Residual U-Net, which consists of two main components: the U-Net architecture itself (Fig. 7) with an encoder that downsamples the image features at multiple scales and a decoder that upsamples these features to produce a segmentation map with the same resolution as the input. Each block of the encoder and the decoder is connected by skip connections, allowing the decoder to access information from different scales. Both the encoder and decoder consist of a 5 layer network with, respectively, downsampling and upsampling by a factor of 2 at each layer. The Residual U-Net combines the concepts of residual networks (ResNets) and the U-Net architecture such that each layer in the encoder and decoder is implemented as a residual block (Fig. 8) which contains skip connections that allow the network to learn residual information: each block consists of a residual unit with 2 sub-units. Each sub-unit has a batch normalization, rectified linear unit (ReLU) activation function and convolution layers with (2,2) kernel filters for each convolution. An additional 20% dropout was applied.

**Figure 8.**
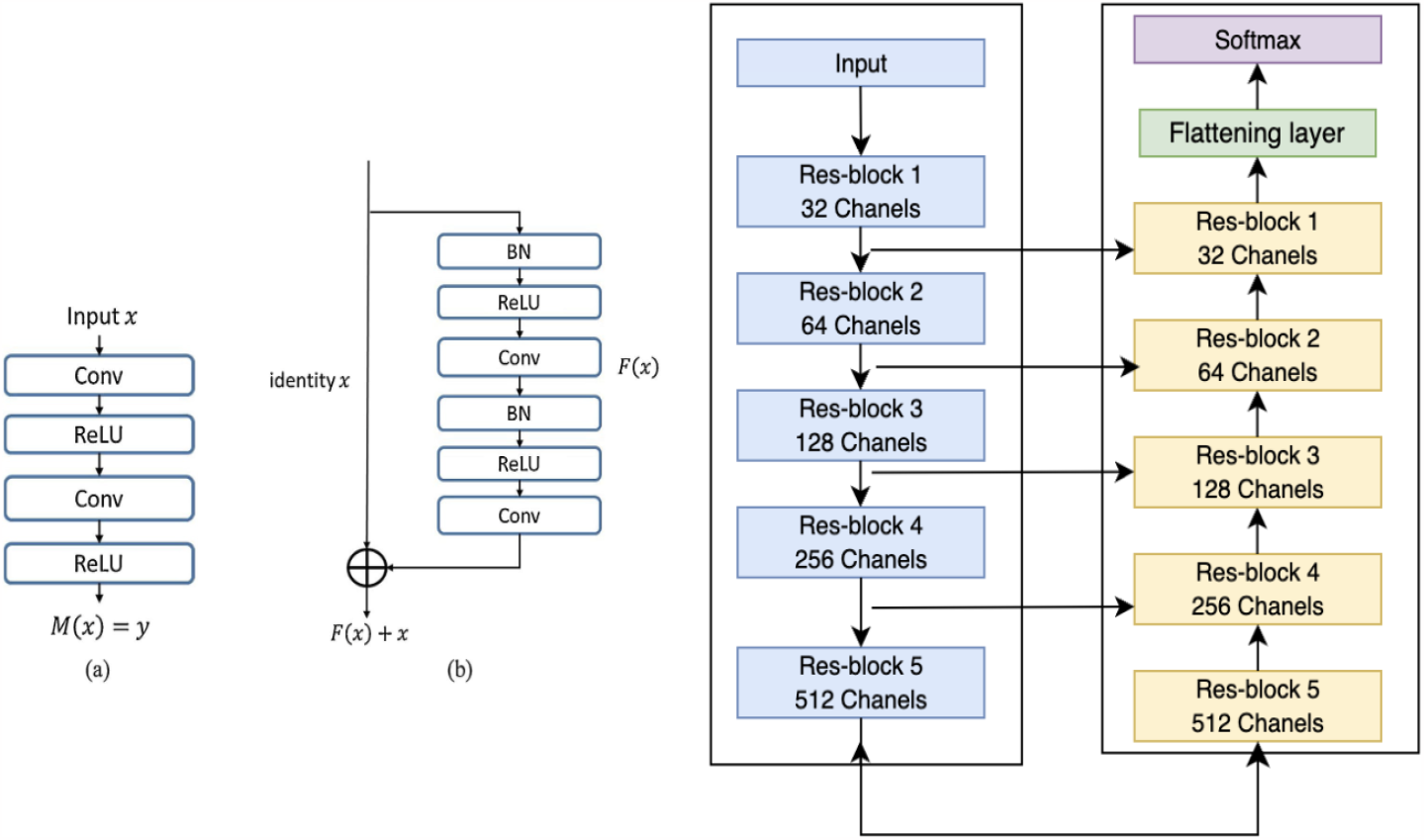
Building blocks of the Residual U-Net architecture with (left image) in (a), plain neural unit used in a standard U-Net and (b), a residual unit with identity mapping and (right image) the full U-Net architecture with Encoder-Decoder components.

The output of the decoder is a logit vector, we then use the Softmax function given that we are classifying into 3 separate classes and thus the loss function used is the weighted cross-entropy loss. Indeed, as we mentioned previously, the data set is highly imbalanced, which is why it was necessary to add the class weights in the loss function. The evaluation metrics considered here are the precision, recall, accuracy and IOU (Intersection Over Union) coefficient.

The Softmax function, cross-entropy and balanced cross-entropy losses (Equations (1)) are defined as follows:

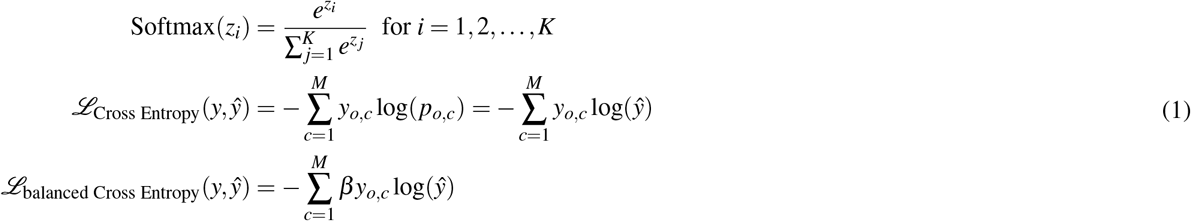

where M is the number of classes (in this case 3), log is the natural log, y is the binary indicator (0 or 1) if class label c is the correct classification for observation o and p is the predicted probability for observation o being in of class c (we can also write 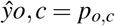. In the balanced cross-entropy, *β* represents the weight of each class and where *β* ∈ [0, 1]. We also define the IOU (Intersection Over Union) (Equation (2)) the following way:

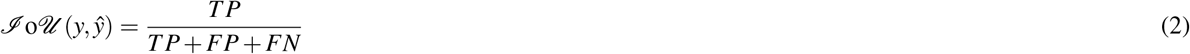

### 3.8 Data post-processing and final predictions

The proposed algorithm improves segmentation results by also exploring the EcoFab substrate geometry to define the key region-of-interest (ROI). It combines prior knowledge about plant root evolution in these specialized hydroponic systems. To minimize the noise surrounding the segmented root, we employed morphological operations^27,28^, starting with dilation on the stacked segmented root image obtained from predictions on all EcoFab acquisition dates. We then extracted a convex hull from the segmented roots and applied area closing and area opening transformations.

To calculate the convex hull of a set of points, we used the Graham’s Scan algorithm, which main steps are as follows:

- Identify a pivot point, usually the point with the lowest y-coordinate (and leftmost if tied).
- Sort all points by their polar angles with respect to the pivot.
- Initialize an empty stack to store convex hull points.
- Iterate through the sorted points and add them to the stack while ensuring a counterclockwise order.
- The stack’s content represents the convex hull vertices.

We used Python libraries such as ‘scipy’ to efficiently compute the convex hull, which finds the smallest convex polygon enclosing all the given points.

Utilizing a convex hull for ROI delimination consists of employing a geometric shape that encompasses a set of points while forming a convex polygon with the minimum possible area. In this context, the points represent the set of stacked roots (maximum intensity projection of timeseries prediction) for a specific EcoFab, determined by the coordinates of the root pixels. The convex hull algorithm determines the outermost boundary that encloses all the points, forming a convex shape. By comparing the convex hull with the original stacked noisy image, we were able to identify and remove noise located outside the hull. Through the combination of the convex hull method and morphological operations, we effectively eliminated the noise surrounding the root at each time point (Fig. 9).

**Figure 9.**
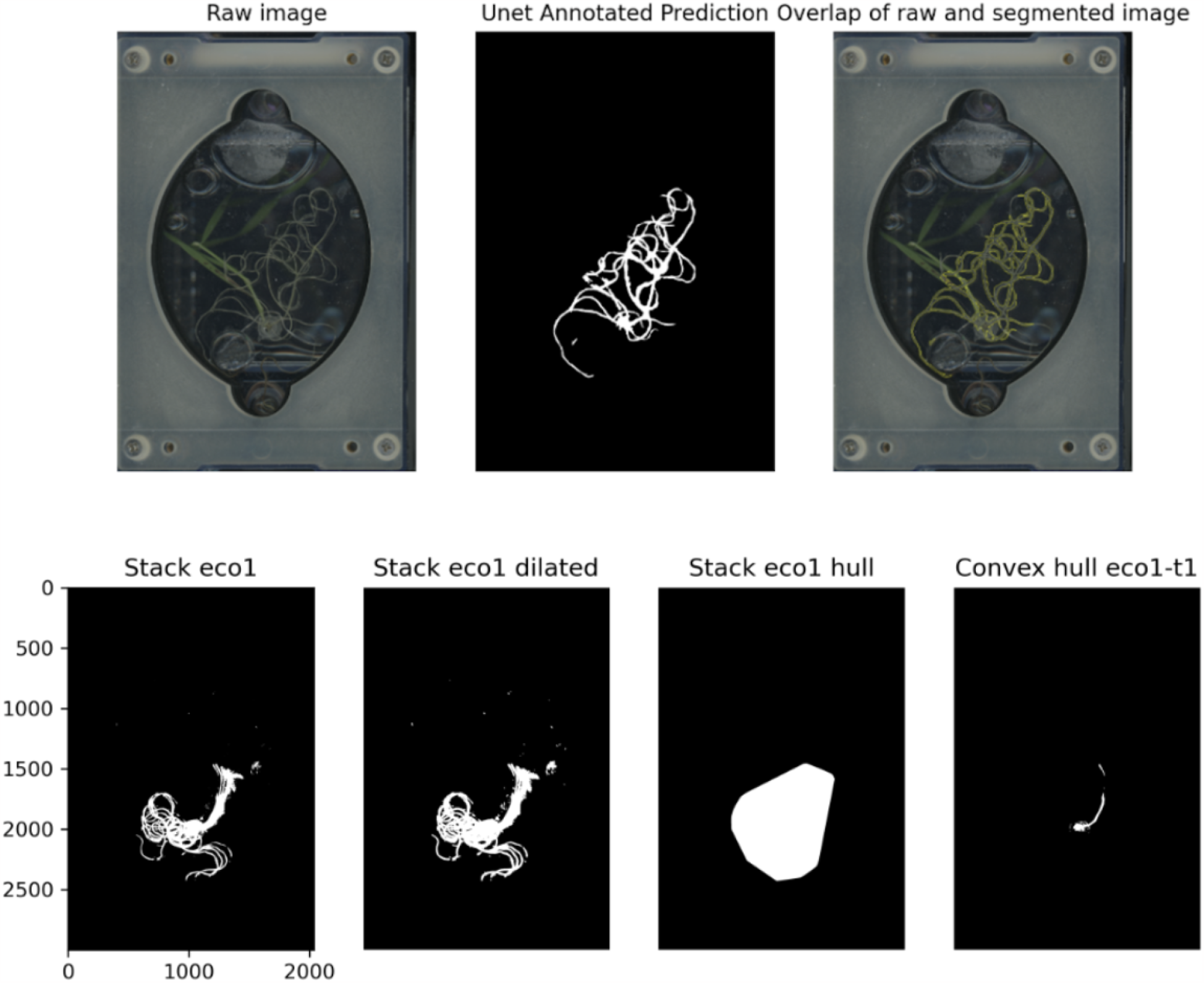
Top: raw, prediction and overlap of segmentation with raw image. Bottom: stack of all prediction masks for a given EcoFab, morphological dilation applied to the stacked result, convex hull applied to the stacked result and final prediction after removal of all surrounding noise

**Figure 10.**
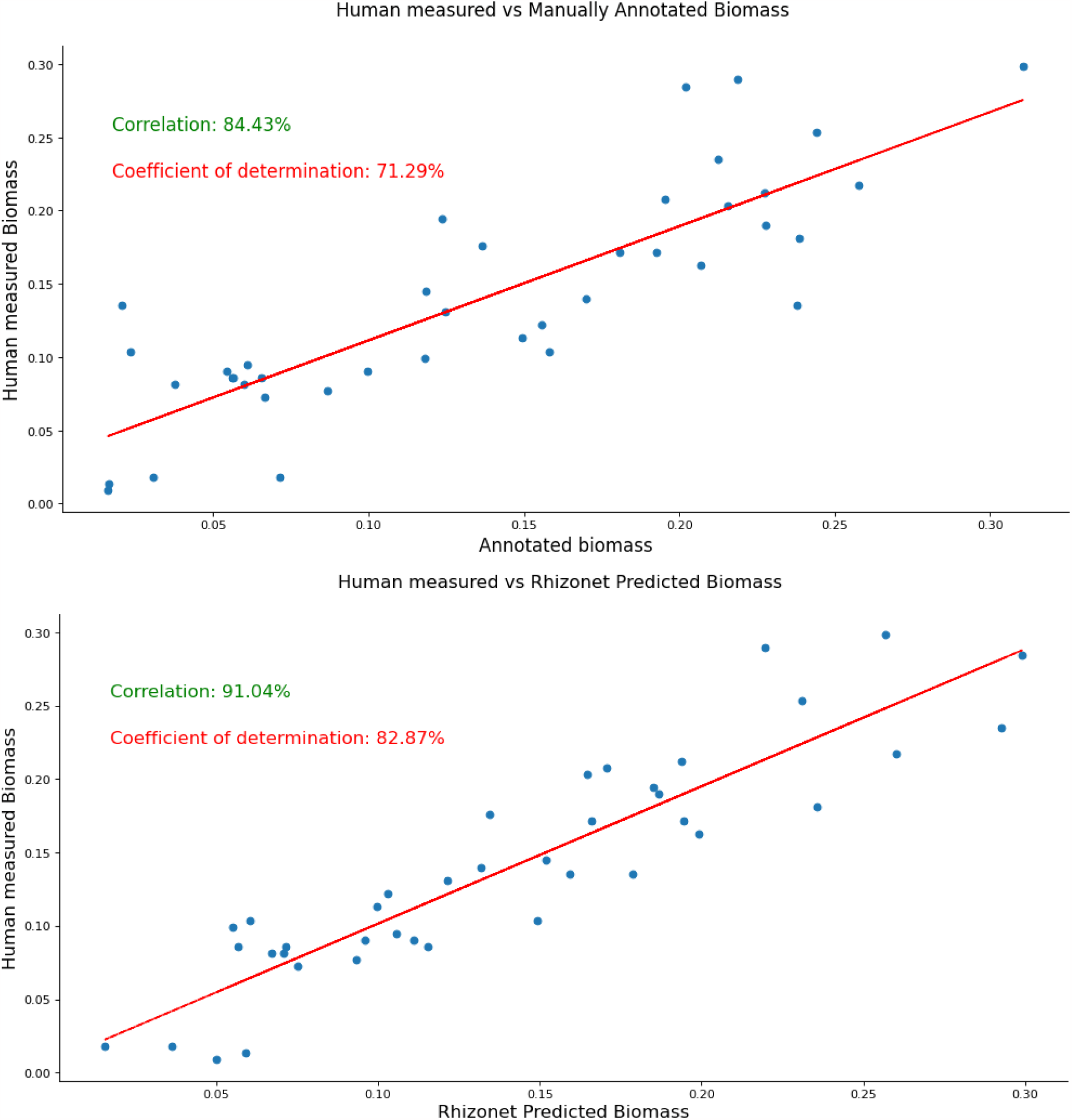
Linear regression of root weight measured by human with scale vs. manual annotation (top) and RhizoNet-predicted biomass (bottom).

## 4 Results

In the context of root segmentation within EcoFAB images, it is essential to consider the challenges posed by physical phenomena such as noise, bubbles, droplets, and shadows. These issues are particularly pronounced at a lower spatial scales, both pixel-wise and during manual annotation. It is reasonable to infer that using smaller patch samples (refer to Table 3) during training helped the algorithm distinguish roots from noise more effectively. Table 4 highlights this last aspect since each metric increases as the patch size decreases.

**Table 4.** RhizoNet without postprocessing for three patch size with performance based on Accuracy, Precision, Recall and IOU.

| Patch Size | Accuracy | Precision | Recall | IOU |
| --- | --- | --- | --- | --- |
| (64,64) | $71.3 \pm 0.58$ | $72.3 \pm 0.58$ | $96.3 \pm 1.53$ | **70** |
| (128,128) | 57 | $56.3 \pm 0.58$ | $96 \pm 0.9$ | $55 \pm 0.85$ |
| (256,256) | $50.3 \pm 1.53$ | $45 \pm 1.73$ | $88 \pm 1.0$ | $43 \pm 1.73$ |

To evaluate the predictions, Fig. 10 (top) shows the comparison between the predicted root biomass with the ground truth of the last acquisition date for each type of media type. We performed a linear regression analysis between predicted biomass and the root weight measured manually using a scale, yielding a correlation value of 71.29% and a coefficient of determination of 84.43%. However, some outliers persisted as a result of the diverse range of artifacts around and over roots, which can closely resemble roots in terms of size and color.

We also assessed the performance of manual annotations in comparison to automated segmentation concerning root weight measured by a human using a balance. As in Fig. 10 (bottom), a correlation value of 91.04% and a coefficient of determination of 82.87% was observed. This improvement in automated segmentation over manual annotations, also observed in Fig.11, underscores the challenge human annotators face when identifying pixels corresponding to roots versus noisy artifacts, especially when these elements share similar thin structures and colors. The algorithm, working with small-size patches and analyzing root structures pixel-wise, appears to be better equipped for this task. In particular, two specific plants with potassium and calcium deficiencies died before the last date in Experiment 2, probably due to lower temperature settings in the growth room. In contrast, most of the 40 EcoFabs in Experiment 1 reached the last date, with the majority exhibiting positive growth.

**Figure 11.**
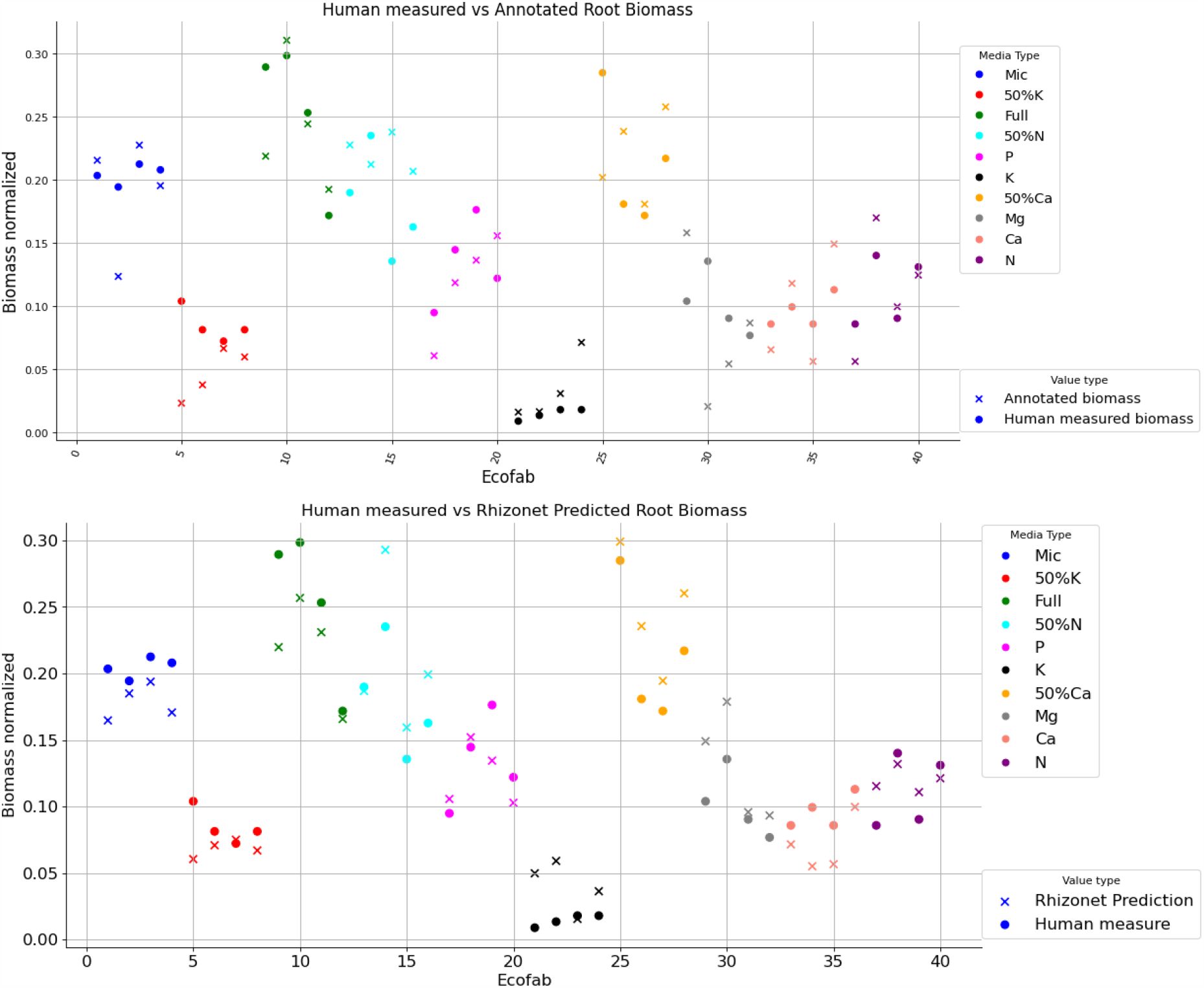
Relationship between root weight measured by human with a scale vs. manual annotation (top) as well as the RhizoNet-predicted biomass (bottom): colors mean different media type or the nutrient condition for each plant.

This highlights the critical influence of environmental factors, including temperature and nutrient availability, during plant growth.

## 5 Conclusions

The Rhizonet model exhibits significant promise for root segmentation, despite the inherent challenges posed by bubbles, condensation, and leaf shadows during root plant data acquisition within the EcoFAB environment. These challenges were effectively addressed through the utilization of a Residual U-Net architecture, enabling the model to discern and separate root plants from unwanted noise in most instances, resulting in precise and accurate segmentation. However, it is important to note that when condensation dominates a large portion of the image, segmentation accuracy may be compromised.

Through a comparative analysis with manual annotations, RhizoNet showcased strong performance metrics, particularly in terms of Intersection over Union (IOU) and accuracy. Incorporation of residual connections in the U-Net architecture proved beneficial, facilitating information flow across network layers. The additional postprocessing operators using convex-hull also enhanced the model’s ability to focus on the best ROI, thereby improving overall performance.

Furthermore, the results of the two experiments underscore the critical role of environmental parameters during plant growth, especially in relation to specific nutrient concentration. This suggests that refining these parameters could enhance project results. Additionally, exploring smaller patch sizes is a promising avenue, since the best results were achieved with a patch size of 64, the smallest among the three tested sizes. Smaller sizes may offer improved feature encoding capabilities within the model.

The unlabeled data played a crucial role in extending the utility of our model. By utilizing these data to make additional predictions with the weights of the trained model, we were able to visually assess its performance on the data not used during training. This evaluation not only demonstrated the model’s capacity for generalization but also showcased its potential for transfer learning.

## Acknowledgements

This work was supported by the US Department of Energy (DOE) Office of Science Advanced Scientific Computing Research (ASCR) and Basic Energy Sciences (BES) under Contract No. DE-AC02-05CH11231 to the Center for Advanced Mathematics for Energy Research Applications (CAMERA) program. It also included support from the DOE ASCR-funded project Analysis and Machine Learning Across Domains (AMLXD) and DOE Biological and Environmental Research (BER), Genomic Science Program funded project Twin Ecosystems Project: A New Capability for Field and Laboratory Ecosystems Coupled by Sensor Networks and Autonomous Controls.

## Author contributions

T.N. and J.S. formulated the project and led research planning, execution, and coordination. P.A. and T.N. conceived the plant experiment(s), P.A. and T.N. conducted the plant experiment(s), P.A., D.U. and Z.S. annotated images, D.U. designed the computer vision and machine learning algorithms in RhizoNet, D.U. and Z.S. developed the computational methods and analyzed the results. All authors contributed to the writing of the article and reviewed the manuscript.

## Additional information

Correspondence and material requests should be addressed to D.U.

